# Myogenic dedifferentiation involves a p53-dependent reorganization of PLK4 localization during centrosome regeneration

**DOI:** 10.1101/2024.06.17.599282

**Authors:** Elaiyaraja Subramanian, Gonçalo Brito, Anoop Kumar, Matthew Kirkham, András Simon

## Abstract

Multinucleated skeletal muscle cells are stably withdrawn from the cell cycle in most vertebrates. Muscle dedifferentiation is however naturally occurring during limb regeneration in newts and can be artificially induced in mammalian myotubes. Dedifferentiation involves fragmentation of myofibers and myotubes into mononucleate cells which subsequently reenter the cell cycle, and give rise to proliferative progeny. Here we addressed the dynamics of centrosomes, which are key organelles for cell proliferation during myogenic differentiation and dedifferentiation. We show that, in contrast to their mammalian counterparts, newt muscle cells retain centrosomes during differentiation and demonstrate that regeneration of centrosomes in dedifferentiated mouse muscle cells depends on inhibition of the tumor suppressor p53. We also find that regulation of the subcellular localization of Polo-Like Kinase 4 rather than its expression level is a hallmark of myogenic differentiation and dedifferentiation, identifying a novel cellular process underlying the plasticity of the differentiated state.

## Introduction

Skeletal muscle differentiation involves stable withdrawal from the cell cycle, which is generally referred to as the postmitotic state (Berkes and Tapscott, 2005; Braun and Gautel, 2011; Walsh and Perlman, 1997). Reversal of myogenic differentiation occurs naturally during limb regeneration and is characterized by three main phases: (i) fragmentation of the syncytium (ii) temporal downregulation of a select set of muscle differentiation markers in the myofiber-derived mononucleate progeny, (iii) which subsequently reenter the cell cycle and proliferate (Kumar et al., 2000; Subramanian et al., 2023; Wang et al., 2015; Wang and Simon, 2016). While muscle dedifferentiation in newts has been well described at the cellular level (Calve and Simon, 2011; Hay, 1959) and some of its inductive molecular underpinnings identified (Sandoval-Guzmán et al., 2014; Subramanian et al., 2023; Tanaka et al., 1997; Velloso et al., 2000; Wang et al., 2015), the dynamics of centrosomes with microtubule organizing capacity, which are key cellular organelles for cell division (Bornens, 2012; Conduit et al., 2015; Fu et al., 2015; Nigg and Raff, 2009; Sir et al., 2013), has remained unexplored. In mammals, centrosomes are lost during muscle differentiation (Connolly et al., 1986; Musa et al., 2003; Przybylski, 1971; Tassin et al., 1985), and microtubule organization shifts to non-centrosomal locations (Becker et al., 2020; Srsen et al., 2009; Tassin et al., 1985). Hence, three principal scenarios are possible in newts: (i) differentiated muscle cells could retain centrosomes in newt; (ii) centrosomes could be lost, similarly to mammals, but generated *de novo* (regenerated) because of or at least concomitant with dedifferentiation; (iii) myofiber-derived mononucleate cells could regain proliferative potential without centrosomes. Although being a rare phenomenon, cell proliferation without centrosomes has been described in certain contexts, like development of fruit flies and mice (Basto et al., 2006; Howe and Fitzharris, 2013), and at whole organism level in asexually reproducing flatworms (Azimzadeh et al., 2012).

In contrast to newts, skeletal muscle dedifferentiation does not occur naturally in mammals (Bryson-Richardson and Currie, 2008; Tedesco et al., 2010). Nevertheless, dedifferentiation is possible to induce artificially in cultures of myotubes by myoseverin, which causes microtubule depolymerization and fragmentation of myotubes into mononucleate cells (Rosania et al., 2000). Rigorous cell tracking experiments demonstrated that myoseverin-induced myotube-fragmentation produces non-proliferative mononucleate cells (Duckmanton et al., 2005) because of the initiation of an apoptotic program (Wang et al., 2015). Importantly, inhibition of both caspases (Howe and Fitzharris, 2013) as well as the tumor suppressor, p53, in combination with myoseverin treatment is required for myotube-derived mononucleate cells to become proliferative (Wang et al., 2015). Upon expansion in culture, these cells are able to contribute to muscle regeneration in mice (Wang et. al, 2015)

Here we used this experimentally tractable cell-based assay system to ask if centrosomes become regenerated in the myotube-derived mammalian mononucleate progeny as these cells reenter the cell cycle and reach S-phase. We show that centrosomes become detectable in dedifferentiated cells only if p53 is inhibited. Furthermore, we identify that the expression of a key regulator of centrosome assembly, polo-like kinase 4 (PLK4) (Barr et al., 2004; Bettencourt-Dias et al., 2005) remains stable during muscle differentiation. Instead of change in expression level, PLK4 shifts its subcellular localization during myogenic differentiation and dedifferentiation in a manner that is dependent on centrosomal protein 152 (cep152) expression. We also show that newt muscle cells retain their centrosomes with microtubule-organizing activity in their differentiated state. Hence, the data altogether provide both an explanation for why newt but not mammalian muscle gives rise to proliferating progeny after an injury as well as define a link between p53 and the dynamics of PLK4 subcellular localization in the context of muscle dedifferentiation.

## Results

### Centrosome organization in differentiating muscle cells

We compared mouse and newt myogenic cells which were stimulated to undergo differentiation resulting in the generation of multinucleate myotubes expressing the differentiation marker myosin heavy chain (MHC) (Fig 1A-A’’). These differentiation events are well-documented, and the myotubes attain stable postmitotic arrest in both mouse and newt myotubes (Pajcini et al., 2010; Sandoval-Guzmán et al., 2014; Walsh and Perlman, 1997; Wang and Simon, 2016). While we found that mouse myotubes gradually lost centrosomes, newt myotubes retained centrosomes indicated by canonical γ-tubulin localization (Figure 1A-A’’). Having established that newt myotubes differ from their mammalian counterparts in terms of the presence of centrosomes, we next asked whether these features are retained in newt myofibers freshly isolated from mature limbs (Suppl. figure 1A, B, C). We evaluated the presence of centrosomes in myofibers by analyzing the expression of γ-tubulin and CEP135. We found expression and co-localization of these two markers in typical centrosomal patterns (Figure 1B-B’’), out of which 54% had a classical juxtanuclear position (Figure 1C).

**Figure 1.**
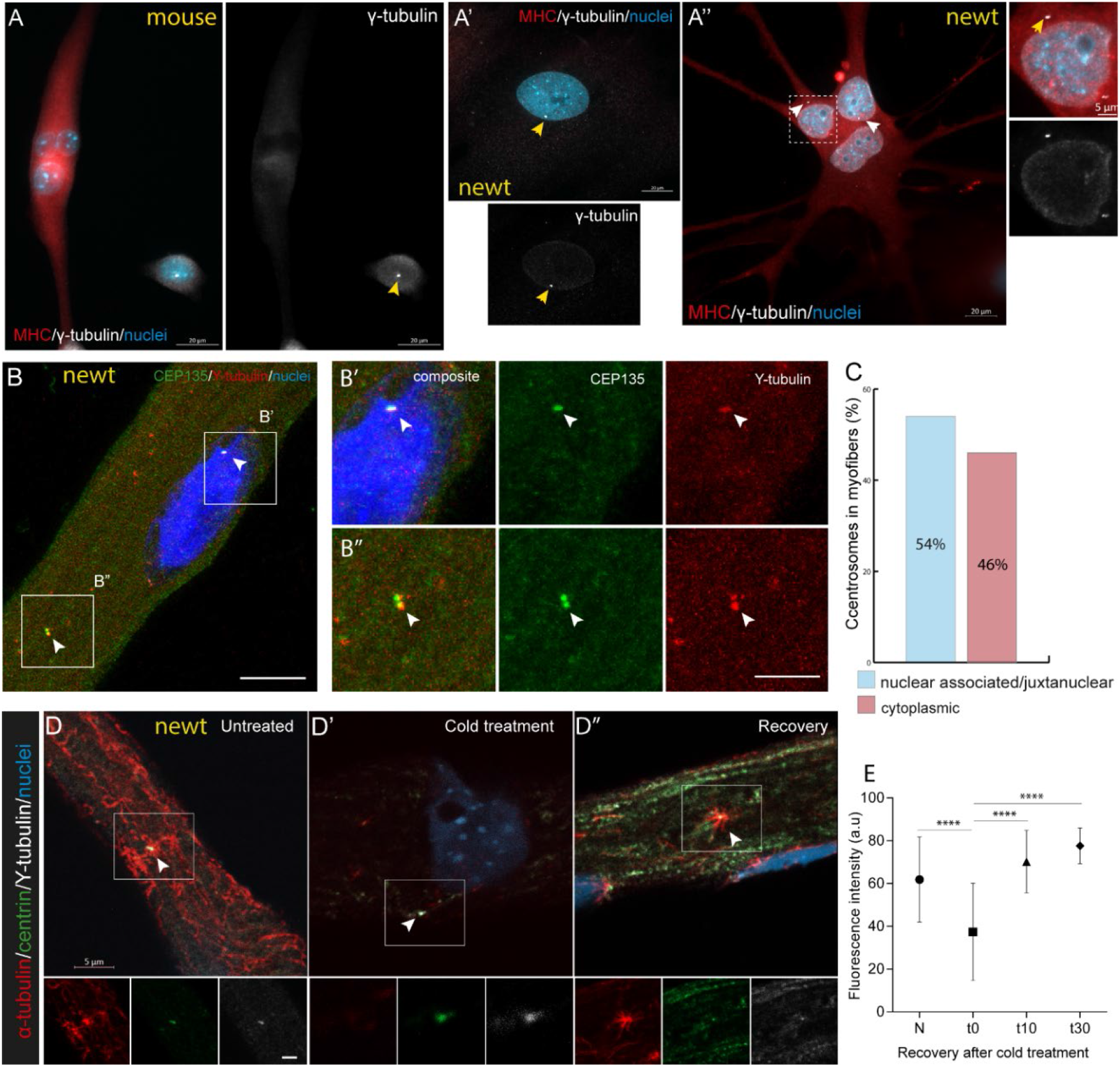
In contrast to mouse muscle, differentiated newt muscle cells retain centrosomes. (**A-A”**) Differentiated mouse and newt muscle cells in culture. (A) A differentiated mouse myotube without centrosomes. Note the centrosome in adjacent residual mononucleate cell (left panel, arrowed yellow). (A’) A newt mononucleate cell with centrosome (arrowed yellow), (A”) Multinucleate newt myotube showing centrosomes (inset, enlarged on the right panel). (**B**) An area of a newt myofiber showing immunostaining of centrosomes with two markers, CEP135 and γ-tubulin. (B’, B’’) Magnified area showing co-immunostaining with the antibodies to the centrosomes (arrowed white) (**C**) Association of centrosomes with nuclei or along the myofiber cytoplasm. Centrosome location varies in myofibers with 54% of the centrosomes being associated with myonuclei (juxtanuclear, B’) and the remaining 46% being located along the myofiber cytoplasm (distant, B’’). A total of 239 centrosomes were analyzed from myofibers pooled from the limbs of 6 different animals. (**D-D’’)** Microtubule regrowth assay. The microtubules were depolymerized through cold treatment on ice for three hours and later allowed to recover at normal temperature. After 10 seconds, recovery and repolymerization of microtubules occurs from the centrosomes. Lower panels show corresponding area magnified from the upper panel. (E) Quantification of mean intensity from α-tubulin during microtubule regrowth assay. The amount of α-tubulin is significantly reduced after cold treatment and rapidly restored upon recovery. N, normal myofibers (N = 63), t0, 3 h after cold treatment (N=63); t10, 10 seconds after recovery (N=99); t30, 30 seconds after recovery (N =69). Data are presented as mean ± SD (****= p< 0.0001). Scale bars, (A-A”) 20 µm, (B) 10 µm, (B’, B’’) 2 µm, (D) 5 µm, inset,2 µm.

Next, we assessed whether centrosomes within myofibers were active by evaluating their microtubule organizing activity (MTOC assay) (Zebrowski et al., 2015). We performed microtubule regrowth assay using cold microtubule depolymerization and allowing their subsequent regrowth for in a short interval at physiological temperature. We found that cold treatment efficiently depleted their microtubules and rapid recovery occurred upon shifting of the treated cells to physiological temperature (Figure 1D-D’’). These microtubules emanated from centrosomes that expressed both centrin and γ-tubulin (Figure 1D’’). To validate the microtubule organizing activity of centrosomes in myofibers, we quantified the intensity of α-tubulin during microtubule regrowth. The intensity was significantly reduced upon cold treatment, and subsequently restored to normalcy at physiological temperature (Figure 1E). Taken together, we conclude that differentiated newt muscle cells differ from their mammalian counterparts by retaining an active state of centrosome components that can act as microtubule organizing centers.

### Centrosomes are retained in dedifferentiated muscle-derived newt limb blastemal cells

Newt myofibers show remarkable cellular plasticity and dedifferentiate to generate proliferating mononucleate cells during limb regeneration. This process has been demonstrated using several lineage tracing strategies including transgenic reporter lines (Kumar et al., 2000; Sandoval-Guzmán et al., 2014; Subramanian et al., 2023). However, whether the resultant mononucleate progeny have centrosomes has not been delineated. Therefore, we used two different newt transgenic reporter lines (loxP-GFP-loxP-mCherry and loxP-mCherry-loxp-H2B:YFP) (Eroglu et al., 2022; Joven et al., 2018; Subramanian et al., 2023) to evaluate the possible inheritance of centrosomes in myofiber-derived cells. Upon muscle-specific Cre expression and resultant recombination, myofibers and their progeny cells become permanently labelled with mCherry or nuclear YFP, depending on which reporter line is used. A schematic of the experimental procedures is outlined in Figure 2A and Suppl fig 2A, 3A. Cre was induced by focal electroporation into the right limb, which resulted in recombination, while no recombination events were observed in contralateral control limb (Suppl. figure 2B, C). The nucleotide analogue, EdU, was used to capture the cell cycle reentry by myofiber-derived progeny cells during blastema formation. Mid-bud limb blastemas were collected and serial longitudinal sections were processed for immunofluorescence. We detected mCherry+/EdU+ cells with centrosomes as indicated by γ-tubulin staining (Figure 2B, C). To corroborate that myofiber-derived cells contained centrosomes, we used the cre-inducible reporter line (loxP-mCherry-loxp-H2B:YFP) which resulted in the appearance of nYFP+/EdU+/ γ-tubulin+ cells in the limb blastema (Suppl figure 3B, C). Therefore, we conclude that dedifferentiated myofiber-derived mononucleate cells form a proliferative population containing centrosomal structures.

**Figure 2.**
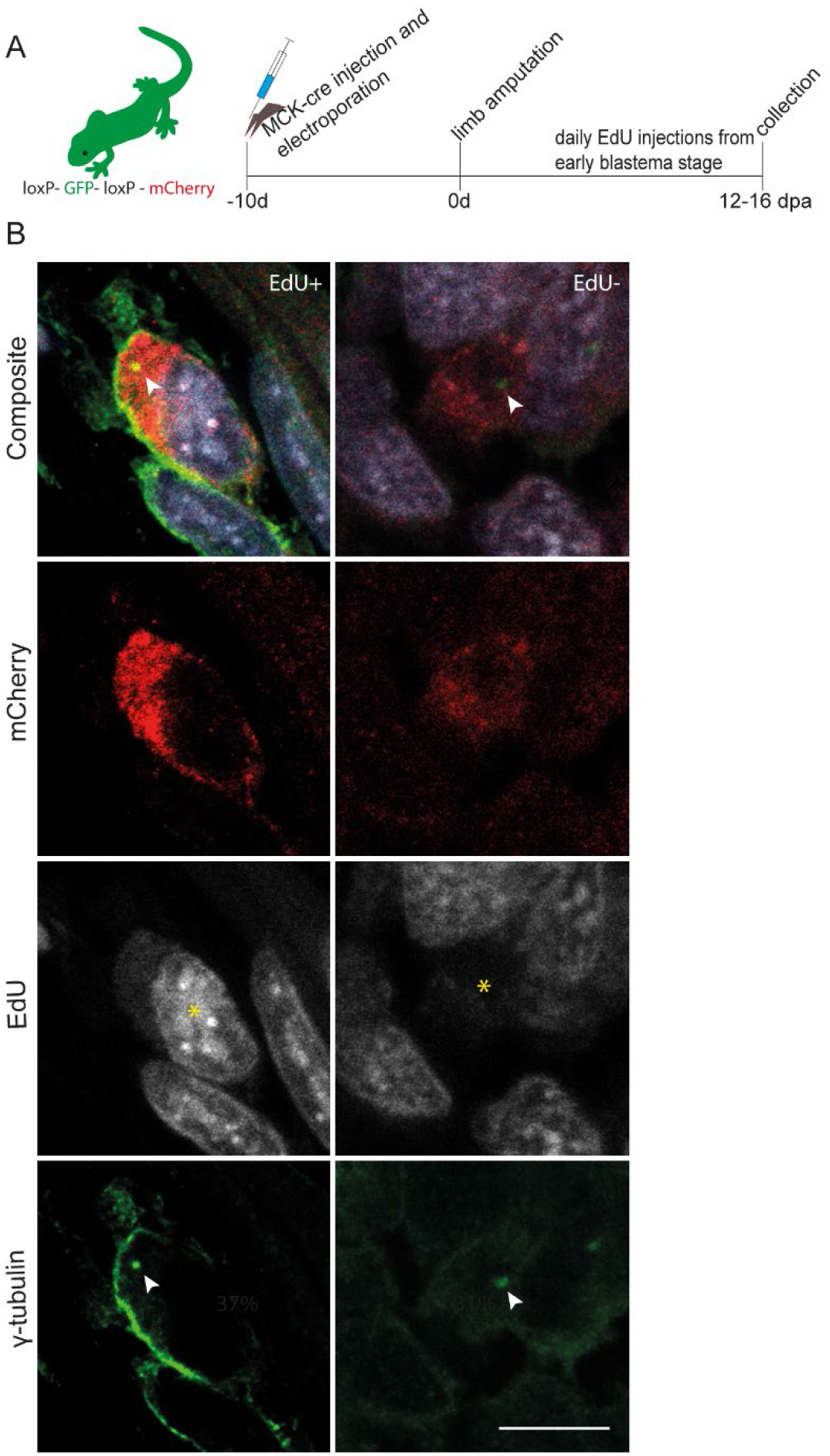
Centrosomes are retained by dedifferentiated mononucleate cells during newt limb regeneration. (**A**) Outline of the experiment illustrating Cre-induced recombination leading to mCherry expression in newt limbs. (**B**) Immunostaining on newt limb sections showing γ-tubulin-positive centrosomes (arrowed white) in a dedifferentiated mononucleate cell (mCherry+ /EdU+), and non-proliferating dedifferentiated muscle cells (mCherry^+^/EdU-; right panel). Yellow asterisk indicates EdU+ or EdU-cells in respective panels. Scale bar, 10 μm.

### Regeneration of centrosomes during dedifferentiation of mouse myotubes

To test whether forced dedifferentiation of mouse myotubes leads to regeneration of centrosomes in the mononucleate progeny, we optimized the protocol by Wang, 2015 (Wang et al., 2015). A schematic drawing of the assay protocol is shown in Figure 3A, (for details, see Methods). In brief, cellularization of myotubes was induced by myoseverin-treatment and subsequent cell cycle reentry of the resultant mononucleate, myotube-derived cells were achieved by inhibition of p53 using pifithrin-a. For specifically tracing myotube-derived cells, we fused primary myoblasts carrying a reporter which encodes for conditional expression of tdTomato with myoblasts expressing cre-recombinase under the control of the muscle creatine kinase (MCK) promoter.

**Figure 3.**
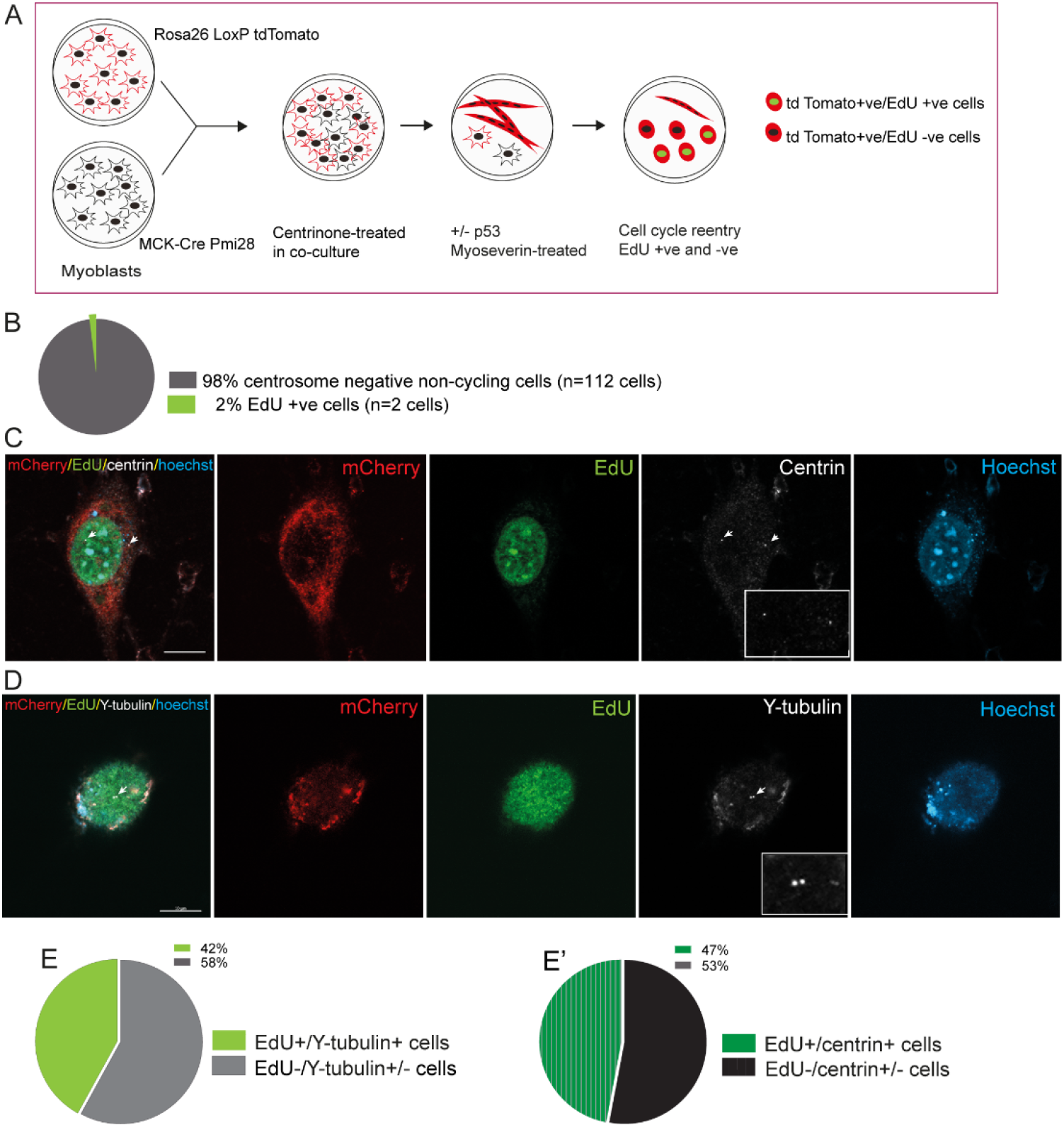
p53-inhibition dependent regeneration of centrosomes during mouse muscle dedifferentiation. (**A**) Schematic showing the assay protocol. Rosa26 myoblasts (CAG:loxP-stop-loxP:tdTomato) and Pmi28 myoblasts (MCK-Cre) were cultured separately in presence of centrinone (5 µM) for four days. The cells were plated together at 3:1 ratio in differentiation medium in presence of centrinone. During the fusion stage, on day 5 the cultures were treated with p53 inhibitor (Pifithrin-α, 60 µg/mL) for 24 h. Cellularisation was induced by treatment with myoseverin (25 µM) for 24 h. Thereafter, the medium was changed to growth medium containing PCD inhibitors and growth factors, and EdU was incorporated to measure cell cycle reentry. (**B)** Distribution of mononucleate cells in myotube cellularisation assay without p53 inhibitor. (**C**) A dedifferentiated mononucleate cell showing cell cycle reentry and immunoreactivity with centrin. (**D**) A dedifferentiated cell showing cell cycle reentry and immunoreactivity with Ƴ-tubulin. (**E, E’**) Distribution of centrosome markers and S phase entry in dedifferentiated myotube-derived mononucleate cells in presence of p53 inhibitor. Scale bar, 10 μm; inset, 0.5 μm.

To ensure that no residual centrosomes were present in the myotubes before the dedifferentiation assay, we treated myoblasts with centrinone, which eliminates centrosomes (Ng et al., 2020; Wong et al., 2015). Based on γ-tubulin staining, 99.4% of the myotubes thus formed lacked centrosomes in the cultures (Suppl. figure 4). In accordance with previous data (Wang, 2015), cellularization upon myoseverin treatment occurred irrespective of p53 inhibition. Myoseverin treatment without p53 inhibition resulted in generation of non-proliferative tdTomato+ mononucleate cells as determined by EdU incorporation (n=112/114); Figure 3B).

To evaluate the possible regeneration of centrosomes in myotube-derived, mononucleate progeny, we analyzed the expression of two different centrosome markers, centrin and γ-tubulin. These analyses revealed myotube-derived tdTomato+/EdU+/centrin+ as well as tdTomato+/EdU+/ γ-tubulin+ cells, indicating the regeneration of centrosomes following myogenic dedifferentiation (Figure 3 C, D). In independent, parallel assays, we found that 42% γ-tubulin+ and 53% centrin+ cells had incorporated EdU (Figure 3E, E’). A substantial portion of centrosome containing cells were EdU-(Suppl. figure 5), similar to the observations in the differentiated mononucleate cells in the newt limb blastema. This population may represent cells in which S-phase did not coincide with the EdU pulsing. Since all tdTomato+/EdU+ cells had centrosomes, the data collectively indicate that regeneration of centrosomes is critical to generate proliferative mononucleate progeny in a p53 inhibition-dependent manner during muscle dedifferentiation.

### Dynamic subcellular relocalization of PLK4 during muscle differentiation and dedifferentiation

Disassembly of the centrosomes during muscle differentiation and their reappearance in myotube-derived progeny prompted us to examine the status of PLK4, which is a key regulator or centrosome function (REF), during muscle differentiation and dedifferentiation.

We found that PLK4 was localized to the centrosomes in proliferating mouse myoblasts (Figure 4A-A’’’). Incorporation of EdU indicated that centrosomal localization of PLK4 was not dependent on cell cycle stages in myoblasts, since both EdU+ as well as EdU-myoblasts showed the same localization pattern (Suppl figure 6). Upon differentiation, a shift of PLK4 localization occurred from the centrioles, appearing throughout the cytoplasmic compartment (Figure 4B). Notably, this switch in the distribution of PLK4 was restricted to cells which were undergoing differentiation, as indicated by fusion and expression of myosin heavy chain, but not in the residual, undifferentiated mononucleate cells (Figure 4B, panel Day 5; Suppl. figure 7B). In fully differentiated myotubes, PLK4 distribution extended throughout the cell and showed juxtanuclear as well as myofibrillar association. (Figure 4B; Suppl. figure 7B).

**Figure 4.**
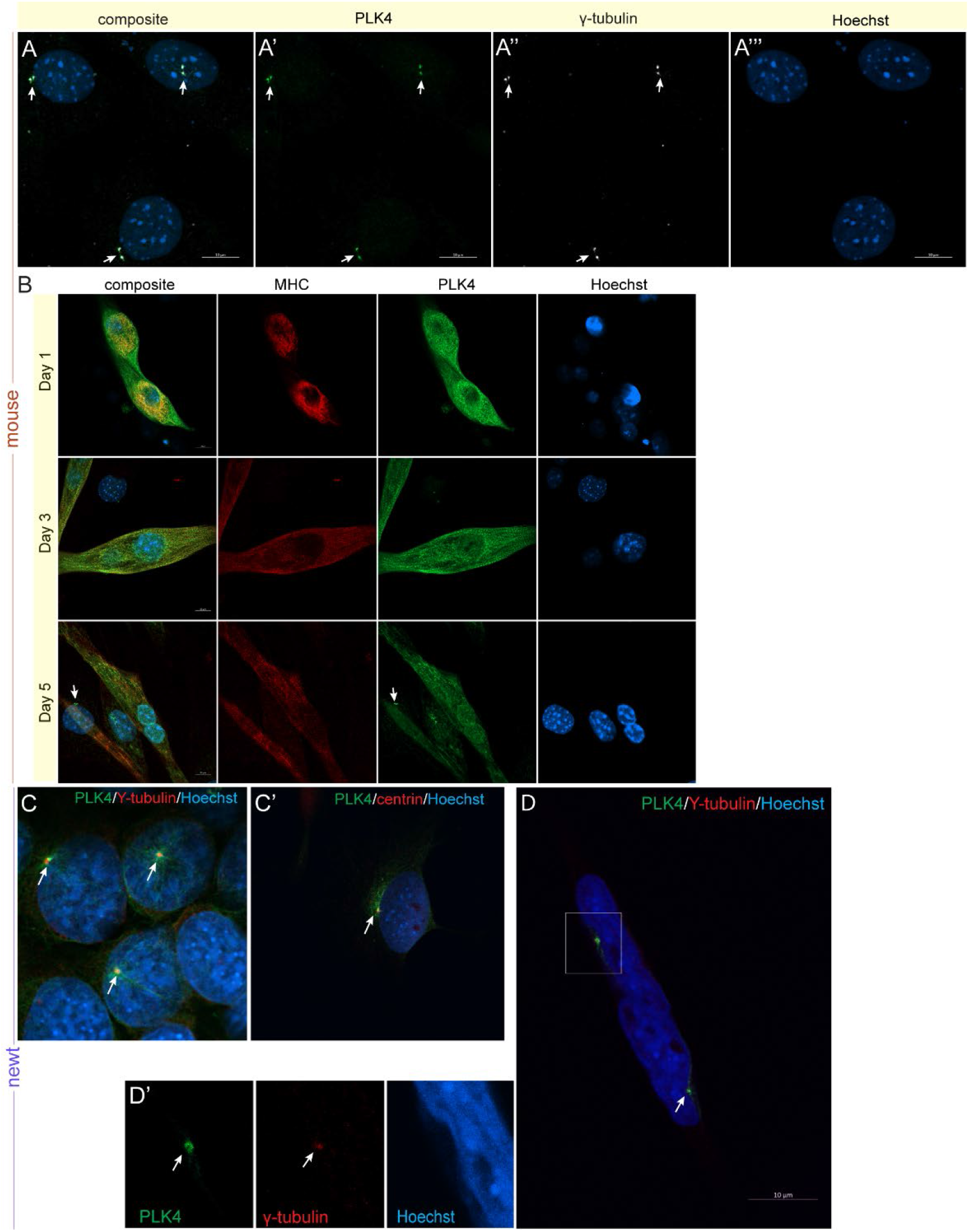
In contrast to newt muscle, PLK4 is redistributed in differentiated mouse muscle cells. (**A-A’”**) PLK4 protein is expressed in mouse myoblasts, and it is localized to centrosomes (arrowed white). (**B**) Progressive differentiation of myoblast to form myotubes in culture. Note the shift in PLK4 from focal centrosome expression to a cytoplasmic pool. By day 3, the differentiating myotubes also show association of PLK4 and sarcomeres. Note the centrosome localization of PLK4 in residual unfused mononucleate cells in day 5 culture (arrowed white). (**C**) PLK4 expression in newt mononucleate cells. PLK4 colocalizes with centrosomal protein γ-tubulin. (**Ć**) PLK4 colocalization with another centrosome marker Centrin in newt cells. (D) Localization of PLK4 in centrosomes of a newt myofiber in culture. Colocalization with γ-tubulin is shown in inset (D’). In all panels, the white arrows point to centrosomes. Scale bars, 10 μm.

As we observed a redistribution of PLK4 during differentiation of mouse myoblasts, we next asked whether a similar shift in protein localization also occurred in newt primary myogenic cells and myofibers. We generated a custom peptide antibody against PLK4 (see Methods). Immunofluorescence assay showed centrosomal colocalization of PLK4, based on both γ-tubulin as well as centrin reactivity (Figure 4C) in mononucleate newt cells.

Importantly, PLK4 was retained within centrosomes and showed colocalization with γ-tubulin (Figure 4D) also in isolated myofibers. Thus, muscle differentiation leads to intracellular reorganization of PLK4 protein in mammalian but not in newt muscle cells.

### Regulation of PLK4 by CEP152 in muscle cells

Recruitment of PLK4 to centrioles requires the centrosomal protein cep152, and a reduction in cep152 expression leads to failure of centriole duplication or centriole loss in various cell types (Cizmecioglu et al., 2010; Kim et al., 2013; Park et al., 2014; Sir et al., 2013). Hence, we next tested whether cep152 regulated PLK4 localization during myogenic differentiation and evaluated their relation during myogenic dedifferentiation.

Differentiated mouse muscle cultures were divided into two pools: a myotube-enriched (90% myotubes) fraction, and a residual myoblast fraction. Western blot analysis showed a significant reduction in cep152 expression in the myotubes compared to myoblasts (Figure 5A, B). In contrast, the PLK4 expression remained unchanged, in line with the observation that PLK4 was redistributed rather than transcriptionally or post-transcriptionally regulated during muscle differentiation (Figure 5A-C; Suppl. figure 8). Immunofluorescence visualization of cep152 confirmed the reduction of cep152 expression in myotubes (Figure 5D-F’).

**Figure 5.**
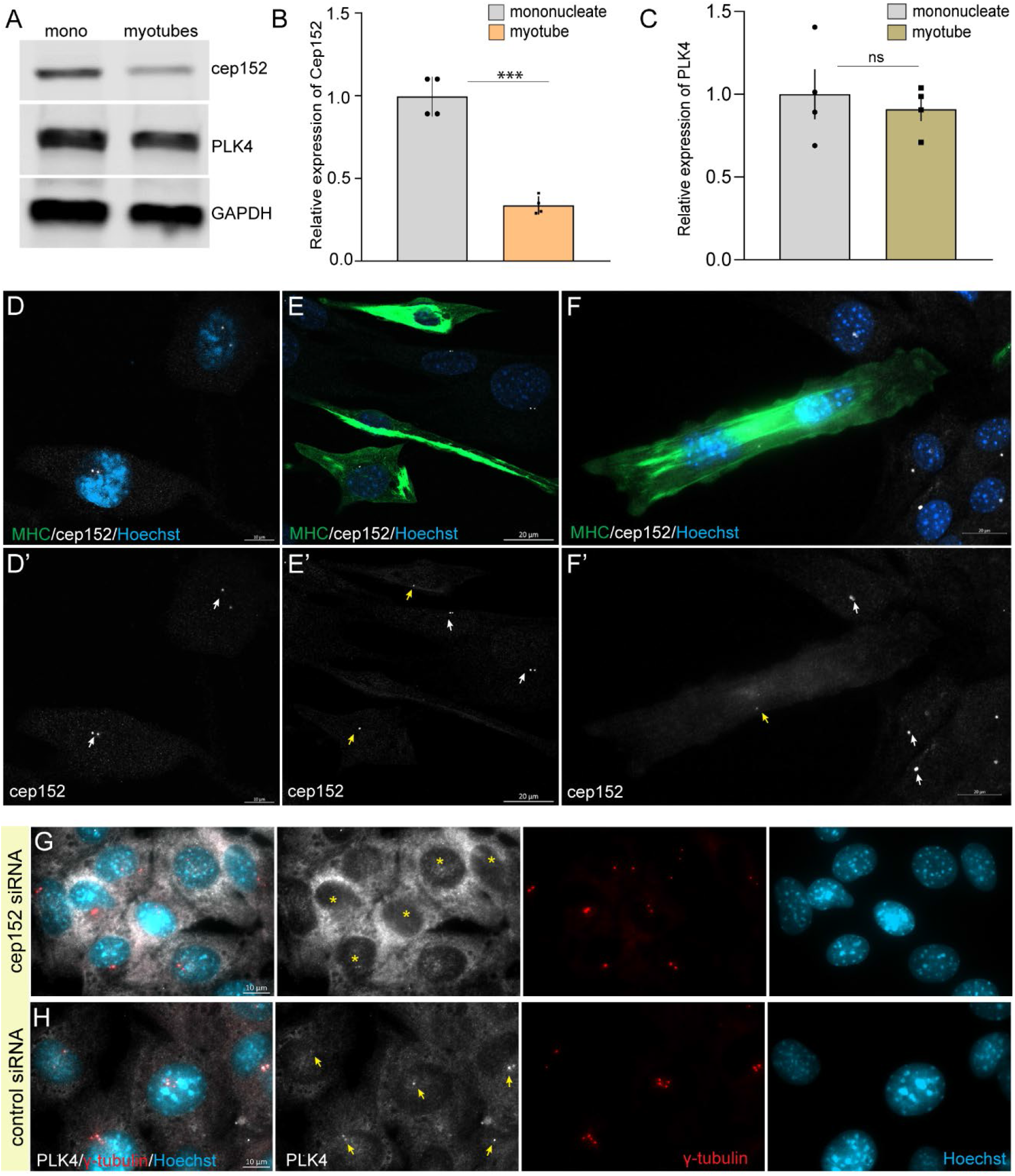
Regulation of cep152 and PLK4 during muscle differentiation. (**A**) Western blot showing cep152, and PLK4 protein in purified fractions of mononucleate cells and myotubes. Note that the blots were run independently for each protein in panel A. (**B**) Relative expression of cep152 in myotube-enriched and mononucleate fractions of pmi28 cells. (**C**) Relative expression of PLK4 in myotube enriched fraction and residual mononucleate pmi28 cells. Note the reduction of cep152 levels in differentiated myotubes. (**D-F**) Progressive loss of cep152 expression during myoblast fusion in culture between days 0-5. (**D’-F’**) The white arrow denotes centrosomes in mononucleate cells whereas yellow arrow indicates weak foci in differentiating cells. (**G, H**) siRNA-mediated knockdown of *cep152* in C2C12 myoblast cells and resultant PLK4 redistribution. (**G**) redistribution of PLK4 protein to cytoplasmic pool (yellow asterisks). (**H**) Expression of PLK4 protein localized o centrosomes in control siRNA-treated cultures (arrowed yellow). In both cases the γ-tubulin remains unchanged. Scale bars, (D-F), 20 μm; (G, H), 10 μm.

To test whether downregulation of cep152 would evoke cytoplasmic distribution of PLK4 without myogenic differentiation, we performed RNAi assay targeted against *cep152*, using three non-overlapping siRNAs. Expression of all three siRNAs resulted in depletion of both *cep152* mRNA and cep152 protein while centrosomal foci were still visible based on y-tubulin staining (Suppl. fig 9). We found that depletion of cep152 resulted in cytoplasmic redistribution of PLK4 (Figure 5G, H), indicating a regulatory role for cep152 in the centrosomal localization of PLK4 in undifferentiated myoblasts.

To determine the relationship between cep152, PLK4, and S-phase reentry during myogenic dedifferentiation, we performed the cellularization assay with or without inhibition of p53 as described in Figure 3A. We observed the presence of centrosomal cep152 in both EdU+ as well as EdU-mononucleate, myotube-derived, dedifferentiated cells (Figure 6A). However, no tdTomato+/ EdU+/CEP152-cells were recorded (N=230). We also found that dedifferentiated tdTomato+ mononucleate cells showed re-localization of PLK4 and formation of centrosomal structures in 70% (N=146) of the cases, out of which 40% were EdU+ and 30% were EdU- (Figure 6B, B’; 6C, C’). The remaining 30% of EdU-cells all showed a cytoplasmic pool of PLK4 which did not relocalize to centrosomes. In addition, we performed the cellularization assay without inhibition of p53 and found that PLK4 failed to relocalize from the cytoplasmic pool when p53 was not inhibited (Suppl. figure 10). In contrast, residual, tdTomato-mononucleate cells, which represent the unfused mononucleate population, maintained PLK4+ centrosomal foci (Suppl. figure 10). Altogether, we conclude that inhibition of p53 is necessary for cep152 regulation and for relocalization of PLK4 to centrosomes during myogenic dedifferentiation.

**Figure 6.**
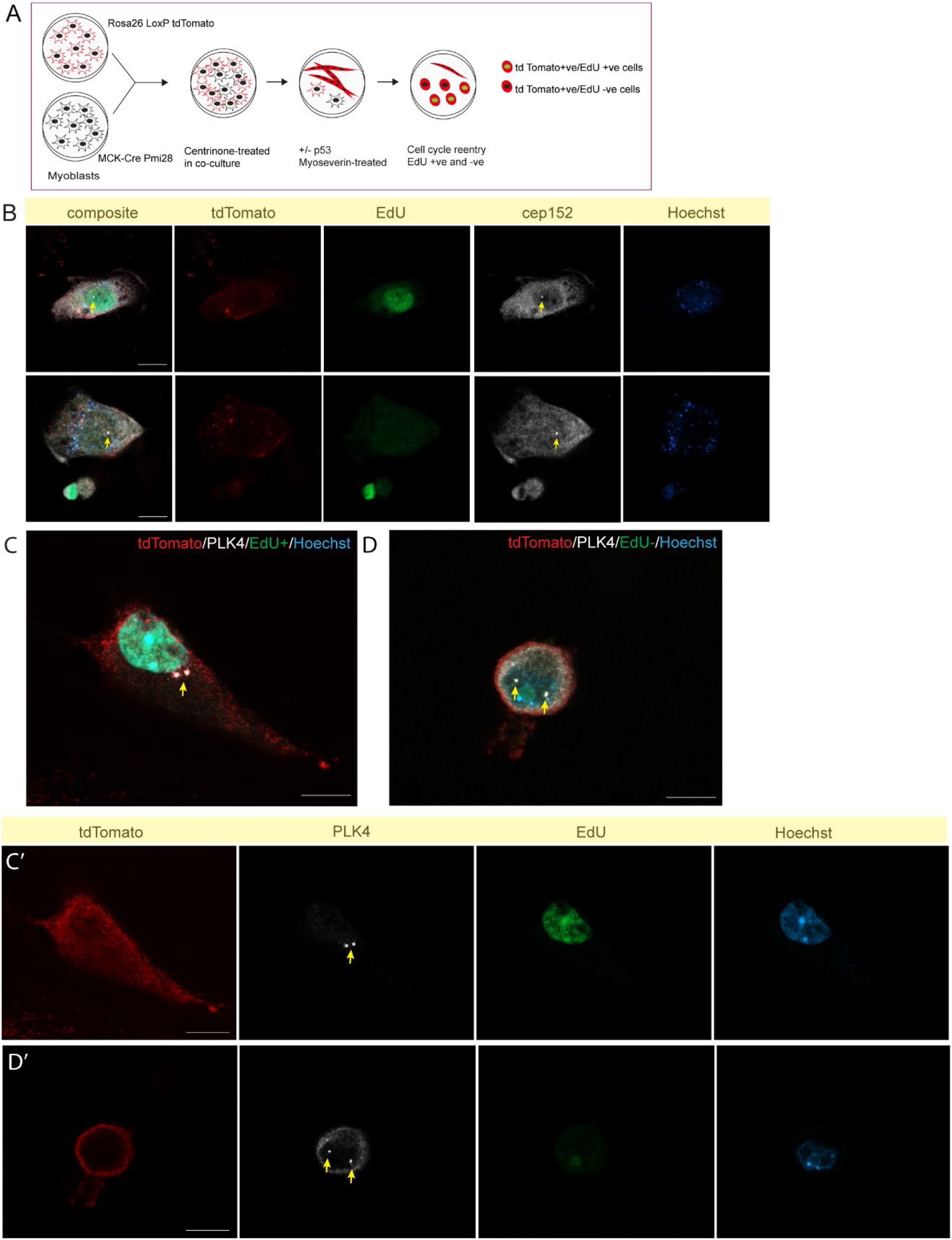
p53-inhibition-dependent regulation of cep152 and PLK4 during cellularization of myotubes. (**A**) Myotube-derived mononucleate cells from cellularization assay. (**B**) Cep152 expression in dedifferentiated cells. Both EdU+ (upper panel) and EdU-(lower panel) mononucleate cells show presence of cep152 protein localized to the centrosome (arrowed yellow). (**C, D**) PLK4 expression in dedifferentiated cells. (**C, C’**) an example showing EdU+/PLK4+ mononucleate cell with centrosome (arrowed yellow). (**D, D’**) an EdU-negative cell showing the presence of PLK4 protein located to the centrioles (arrowed yellow). Scale bars, (A-D), 10 μm.

## Discussion

Centrosomes are critical cellular organelles for proper chromosome segregation during mitosis but there are examples of naturally occurring centrosome-independent cell division events (Azimzadeh et al., 2012; Basto et al., 2006; Howe and Fitzharris, 2013). Here we have shown that fully differentiated newt skeletal myofibers retain centrosomes that act as microtubule organizing centers. This is in sharp contrast to mammalian muscle cells which gradually lose centrosomes during differentiation and microtubule nucleation is organized from non-centrosomal localization (Becker et al., 2020; Bugnard et al., 2005; Duckmanton et al., 2005; Tassin et al., 1985). Thus, retention of centrosomal microtubule organizing activity is a common feature shared between the multinucleate skeletal myofiber and the typically mononucleate cardiomyocyte in newts (Zebrowski et al., 2015). This common denominating feature gives a mechanistic explanation why these two differentiated cell types retain their ability to give rise to proliferating progeny that subsequently contribute to tissue regeneration in newts.

Although a comparable dedifferentiation process does not occur in differentiated mammalian skeletal muscle cells that lost centrosomes naturally, we used here an experimental paradigm in which myotubes are forced into dedifferentiation leading to cell cycle reentry and the generation of proliferative myotube-derived progeny. Since all analyzed EdU+ myotube-derived mononucleate cells had centrosomes, the data indicate that regaining proliferative potential requires centrosome regeneration. Although cell cycle reentry without centrosomes cannot be formally excluded, this may account only for a small fraction of the cases, if for any at all.

Centrosome dynamics has been linked to p53 activity. Increased p53 expression leads to downregulation of PLK4 and consequent failure in centrosome duplication (Bettencourt-Dias et al., 2005; Rosario et al., 2010). Conversely, loss of p53 activity leads to centrosome multiplication (Holland et al., 2012; Nakamura et al., 2013; Vitre et al., 2015). In addition, p53 knockdown allows cells to bypass cell cycle arrest at the G1-phase (Wong et al., 2015). Our results show that cell cycle reentry by dedifferentiated muscle cells required inhibition of p53 activity, and that these EdU-incorporating cells had centrosomes, are in line with these observations. How p53 is linked to centrosome regeneration in dedifferentiated muscle cells remains to be determined but the shift in the subcellular localization of PLK4 gives an entry point to these future investigations.

Given that the expression level of PLK4 has been shown to be downregulated during cell differentiation (Ng et al., 2020), it was unexpected to find that PLK4 remained stable during myogenic differentiation and dedifferentiation. In line with previous observations showing that interaction between cep152 and PLK4 (Cizmecioglu et al., 2010), we found that cep152 was required for centrosomal localization of PLK4. Cep152 levels were downregulated in differentiated myotubes and upregulated upon p53 inhibition during forced dedifferentiation. Thus, a model emerges where retention of PLK4 expression contributes to the capacity of differentiated mammalian muscle cells to dedifferentiate and produce proliferative progeny. In this model, p53-inhibition leads to upregulation of Cep152, which could recruit PLK4 back to centrosomes.

It is noteworthy that salamander limb regeneration involves downregulation of p53 activity, and limb regeneration is inhibited if p53 is stabilized (Yun et al., 2013). Since we found that newt myofibers and their progeny in the blastema retain centrosomes, the function of p53 downregulation during limb regeneration should be other than allowing dedifferentiated muscle cells to proliferate without centrosomes.

Taken together our cross-species comparative cellular assessment identifies a p53-mediated reorganization of centrosome components during cellular dedifferentiation of postmitotic skeletal muscle cells and highlights how the dynamics of centrosome components are linked to the plasticity of the differentiated state.

## Materials and Methods

### Animals and procedures

Iberian newts (*Pleurodeles waltl*) were captive bred in our salamander facility in an environmental-controlled atmosphere according to Swedish animal welfare guidelines. Transgenic lines of *Pleurodeles waltl,* tgSceI(*CAG: loxP-GFP-loxP-Cherry*)^Simon^ and tgTol2(*CAG: loxP-Cherry-loxP-H2B:YFP*)^Simon^ were generated from *Wildtype* newts. All animals were fed with live food or pellets. Larval animals of Stage 50-52 were used for experiments whereas post-metamorphic newts were of age 6-8 months. All surgical procedures were carried out according to approved Swedish ethical permits and post-operative care including analgesics were implemented in caring newts.

### Primary culture of newt myofibers

Primary myofibers were isolated from forelimbs of larval P*. waltl* following the detailed protocol described in (Kumar and Brockes, 2007). The isolated muscle cell cultures were maintained in an incubator at 26 °C with 2.5% CO2 supply.

### Culture of newt cells

Newt cell cultures were derived from the forelimbs of adult *P. waltl* following the general guidelines described (Kumar and Brockes, 2007) with modifications. Briefly, the limb tissues from zeugopodial and stylopodial regions devoid of skin were dissociated in a cocktail of collagenase/dispase solution (Kumar and Brockes, 2007) for 60 min in a shaking water bath at 26 °C, neutralized with serum-free amphibian MEM and filtered through 100 µm filter. The filtrate containing the solution was plated on a gelatin-coated culture plate resulting in attachment and proliferation of cells. These cells were expanded and maintained as newt post metamorphic limb (PML) cells. Cultures from passages 10 or above were used in experiments.

### Culture of mammalian cells

C2C12 myoblasts originated from ATCC cell collections. Pmi28 mouse myoblasts were originally obtained from Anna-Starzinsky-Powitz laboratory (Duckmanton et al., 2005). These cells were cultured on Matrigel-coated (1:10 dilution) flasks. Primary myoblasts were generated from Rosa26 transgenic mouse (*CAG:loxP-stop-loxP:tdTomato*) and obtained through Enric Llorens (Karolinska Institutet). Gastrocnemius muscles from the hindlimbs of a mouse were dissected, minced, and further dissociated in collagenase/dispase solution for one hour. The dissociation solution was neutralized in serum containing medium, filtered through 40 µm filter and the filtrate was pelleted, resuspended in serum containing DMEM and plated on to 60 mm Petri dish. Adherent cells were expanded over generations and passage above five were used in various assays.

### Microtubule regrowth assay

To analyze microtubule regrowth, the isolated myofibers were placed on ice for 3h to induce depolymerization. Subsequently, the myofibers were changed to normal temperature to allow the recovery of the microtubules for 10-30 seconds. Before fixation, myofiber cultures were rinsed for 30 seconds at room temperature with 1% Triton-X100 in PHEM buffer (60 mM PIPES, 25 mM HEPES, 10 mM EGTA, 2 mM MgCl2, pH6.9).

Thereafter the samples were fixed in pre-chilled Methanol at -20°C, permeabilized with 0.5% Triton-X in PBS, blocked with 3% Bovine Serum Albumin (BSA) in PBS for 1 h at room temperature and incubated with the relevant primary antibodies overnight at 4°C. The next day slides were washed with PBS and incubated with the corresponding secondary antibodies for 1 h at room temperature. Antibodies were diluted in blocking buffer and sections were mounted in mounting medium (Dako, #S3023).

### Myotube cellularization assay

Rosa26 myoblasts (CAG:loxP-stop-loxP:tdomato) and Pmi28 myoblasts (MCK-Cre) were cultured separately in presence of centrinone (5 µM) for four days. The cells were trypsinized and plated at 3:1 ratio in a co-culture to induce differentiation in 2% horse serum containing medium (low serum) in presence of centrinone. After two days, centrinone was withdrawn from culture and cells were maintained in low serum medium alone. On day 5, the cultures were treated with a p53 inhibitor, Pifithrin-α (60 µg/mL) for 24 h. On day 6, cellularization was induced in differentiated myotubes by treatment with myoseverin (25 µM) for 24 h. At 18 h during myoseverin treatment, the cultures were supplemented with Pifithrin-α and LIF (1000 U) to sustain p53 inhibition as well as to prevent fusion of dedifferentiated mononucleate cells. At 24 h after myoseverin treatment, the cultures were washed and fresh growth medium containing programmed cell death inhibitors (DIDS, 100 µM; Q-VD 10 µM), growth factors (Pifithrin-α 60 µg/mL LIF, 1000U; EGF 20 µg/mL; bFGF 50 ng/mL) and Thymidine 100 µM (Pajalunga et al., 2017) was added to the culture. In addition, EdU was incorporated (10 µM) to measure S phase entry in cultured cells. After 24h, the cultures were fixed and processed for immunofluorescence.

### Limb regeneration experiments

For the lineage-tracing of dedifferentiated muscle progeny cells, post-metamorphic transgenic newts (CAG:loxp-GFP-STOP-loxp-Cherry) and larval (CAG:loxp-Cherry-stop-loxp-H2B: YFP) transgenic newts were used. The right limbs were injected with a construct containing a muscle-specific promoter muscle creatine kinase (MCK) driving the expression of Cre recombinase (∼4μg/μL), using an Eppendorf FemtoJet 4i pressure injector (#5252000013), while the contralateral limbs were served as un-injected negative controls. Immediately after injection the limbs were electroporated using a Nepa21 Super Electroporator (Nepagene) with two pulses of 70 V for 35ms each (poring pulse), followed by 3 pulses of 50 V for 1s each (transfer pulse). In larval limbs the voltages were two poring pulses of 35V and six transfer pulses of 25V at 50 ms intervals. The expression of the construct induced recombination in the myofibers, leading to the excision of the GFP gene and expression of the mCherry or H2BYFP reporter genes. We found that 10 days are sufficient for recombination events. Forelimbs were amputated at mid-humerus level and allowed to regenerate for 10 days before they were collected. During this time early dedifferentiation occurs from regenerating myofibers and a good proportion of myofiber-derived progeny cells contribute to blastema formation. During early blastema stage (∼10dpa), daily injections of EdU (0.1 mL/newt from a stock of 1mg/mL) were given intraperitoneally until limbs were collected (at 12-16dpa, depending on blastema size). The limbs were collected and fixed overnight in 4% formaldehyde at 4 °C, washed in PBS, cryoprotected in 20% sucrose and embedded in Tissue Tek O.C.T (Sakura) and frozen on dry ice. The samples were sectioned as 14μm longitudinal sections in a NX70 cryostat (Thermofisher).

### Western blotting

Pmi28 myoblast cells were cultured on 10 cm Petri dishes. After 24 h, the growth media was replaced by 2% horse serum containing differentiation media for myoblast fusion and differentiation. On day 5 of differentiation, the myotube and mononucleate cells were separated by filtering the trypsinized cell suspension through a 20µm filter. The protein lysates were prepared from respective cell fraction by using the RIPA lysis buffer. The lysates were processed for Western blotting and after protein transfer, the PVDF membrane (Immobilon cat# IPFL00005) was blocked with the LI-COR Intercept blocking buffer (LI-COR cat # 927-60000). The membrane was incubated overnight in the respective antibody with the GAPDH antibody as control. After rigorous washes, the membrane was incubated with goat anti-rabbit fluorescent IR dye at 680 nm (Biotium cat # 20193-1) and CF 800 goat anti-mouse secondary antibody (Biotium cat# 20831). The membranes were imaged under the LI-COR Odyssey scanner.

### Cep152 RNAi and validation

Mouse C2C12 myoblasts were used for cep152 RNAi experiment using IDT TriFECTa RNAi kit. cep152 specific DsiRNAs to three independent targets (mm.Ri.Cep152.13.1, mm.Ri.Cep152.13.2, mm.Ri.Cep152.13.3) and a negative control DsiRNA (NC1) were obtained from IDT. Respective DsiRNAs were mixed with RNAimax reagent (Thermofisher cat # 13778-150) and reverse transfection was performed by following the standard manufacturer’s protocol. After 4 days of post-transfection, phenotypes of the cells were evaluated in treated cultures. The knockdown effect of all the DsiRNAs was validated by two different assays, (i) mRNA expression of cep152 RNAi through qRT-PCR by using SsoAdvanced Universal SYBR Green Supermix (Bio-Rad cat # 1725271) with the mouse specific primers (Mouse cep152, (5’-3’) F-AAGCAGCATCTCTTGGAGGT; R-TCCTGAGCCTTTTCCATGGT; Mouse Rn18S, (5’-3’) F-GCAATTATTCCCCATGAACG; R-GGCCTCACTAAACCATCCAA). Protein level was assessed by Western blotting as described previously. The blots were assayed by an ECL-based detection method (Thermofisher cat#34579). The images were captured under the Bio-Rad ChemiDoc system using the optimal autoexposure settings.

### Antibodies and immunofluorescence

Myoblast and myotube cultures were routinely fixed in 4% formaldehyde (Merck, cat #) for 10 min at room temperature, washed in PBS and postfixed in 100% methanol at –20 °C for min. Thereafter the samples were briefly air dried and washed in PBS containing 0.5% Triton X-100 (PBST) and blocked in 2.5% BSA. For EdU labelling the samples were processed according to manufacturer’s protocol (Thermofisher). This was followed by antibody incubation of the samples with respective antibodies or in combination overnight at 4 °C. After primary antibody incubation the samples were washed in PBST and incubated with secondary antibodies for one h at room temperature. The samples were washed again, counterstained the nuclei with Hoechst 33258, rinsed in PBS and mounted in Fluromount G.

For Plk4 antibody staining, the samples were fixed in modified cytoskeletal buffer. The buffer solution was prepared by mixing 100mM NaCl, 300 mM sucrose, 3 M MgCl2, 10 mM PIPES in sterile water and the pH adjusted to 6.9. To this solution 1M EGTA and 0.5% Triton X-100 were added just before use. A 4% formaldehyde (from 16% FA: Thermofisher, EM grade) solution was prepared in cytoskeletal buffer to fix cell cultures for Plk4 staining. Before fixation, cultures were quickly rinsed in cytoskeletal buffer and thereafter fixed in the fixative for 15 min at room temperature. Processing for immunofluorescence was as described above.

For EdU labelling, Click-iT Plus EdU Imaging Kit was used throughout (Invitrogen, # C10640), following the manufacturer’s protocol. Cell cultures were incubated with EdU at 10µM overnight in all cases and the EdU labelling was performed before antibody labelling.

### Microscopy and imaging

Muscle cell preparations were imaged under Zeiss LSM 880 or LSM 900 microscope using 63x Plan Apochromatic objective (N.A. 1.4 oil) and acquired using Zeiss ZEN software. In cellularization assays the samples were imaged under a Zeiss Axio Imager Z1 or Axio Observer 7 microscope. The images were captured using Hamamatsu Orca-Flash digital camera integrated to Zeiss ZEN software.

### Statistics

Graphs and statistical analysis were performed in Graphpad Prism (Ver. 9.0). No randomization, sex differences or statistical assumptions were used in designing experiments or analysis. The data were analyzed by Student’s ‘t’ test or non-parametric tests as indicated in the legends at 95% confidence levels. In all cases (n) represents the number of individual replicates in experiments whereas (N) denotes total numbers.

### List of antibodies used in this study

#### Custom antibody to newt Plk4 protein

A custom peptide antibody to two non-overlapping regions of newt Plk4 protein (Peptide-1, ASAIGDKREDFKVLNLLGKGS; Peptide-2, DHRTAYQRDESDCSGRGRKA) was made in host chickens by David’s Biotechnologies, Germany. The reactivity of the antibodies was assessed in cultured newt cells and optimal concentration was determined. Of the two peptides, Peptide-1 gave specific reactivity with the Plk4 protein in newt cells.

#### Commercial Primary Antibodies

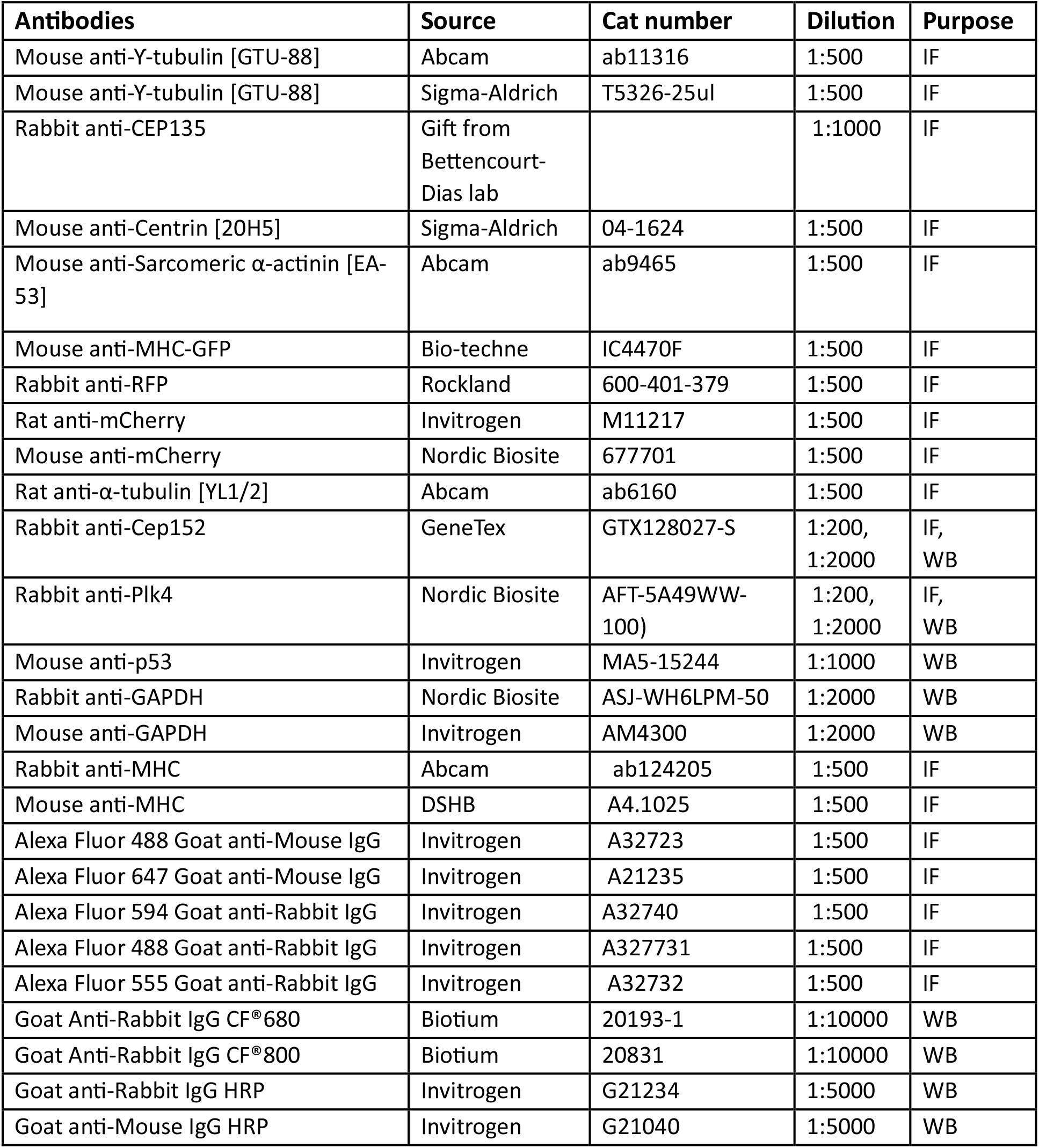

## Acknowledgements

We thank Monica Bettencourt Dias and her lab members for inspiring discussions and sharing unpublished data. We thank Arne Lindquist for critical reading of the manuscript and much helpful suggestions. This work was supported by grants from Vetenskapsrådet, Cancerfonden, and Karolinska Institutet to AS.

---

**Supplementary Figure 1.**
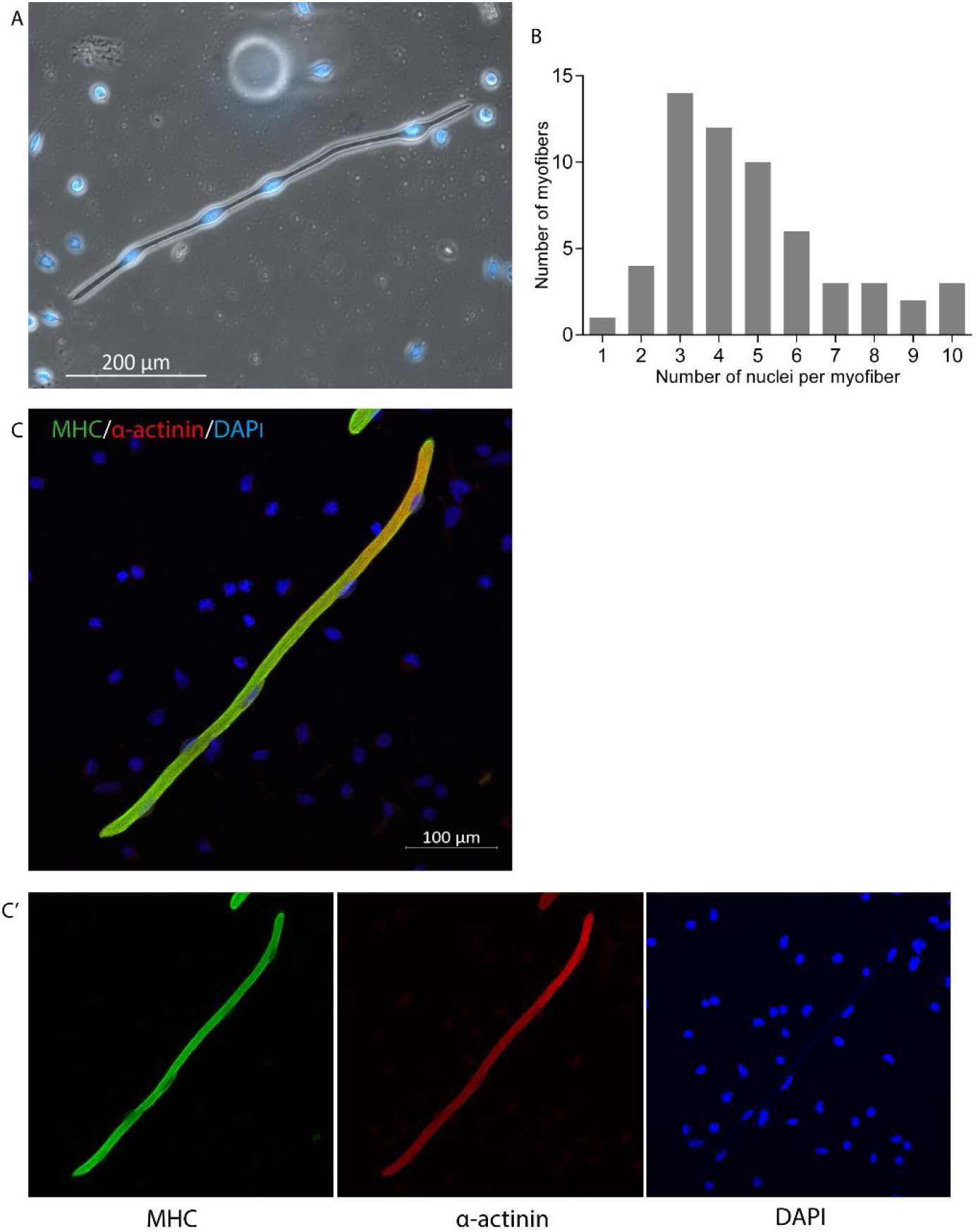
Characterization of myofibers from the Iberian newt, *Pleurodeles waltl*. (**A**) Live myofibers in culture incorporating the nuclear dye Hoechst 33342. (**B**) Distribution of nuclei in myofibers. (**C, C’**) Immunostaining of myofibers with muscle-specific antibodies to Myosin Heavy Chain (MHC) and α-actinin.

**Supplementary figure 2.**
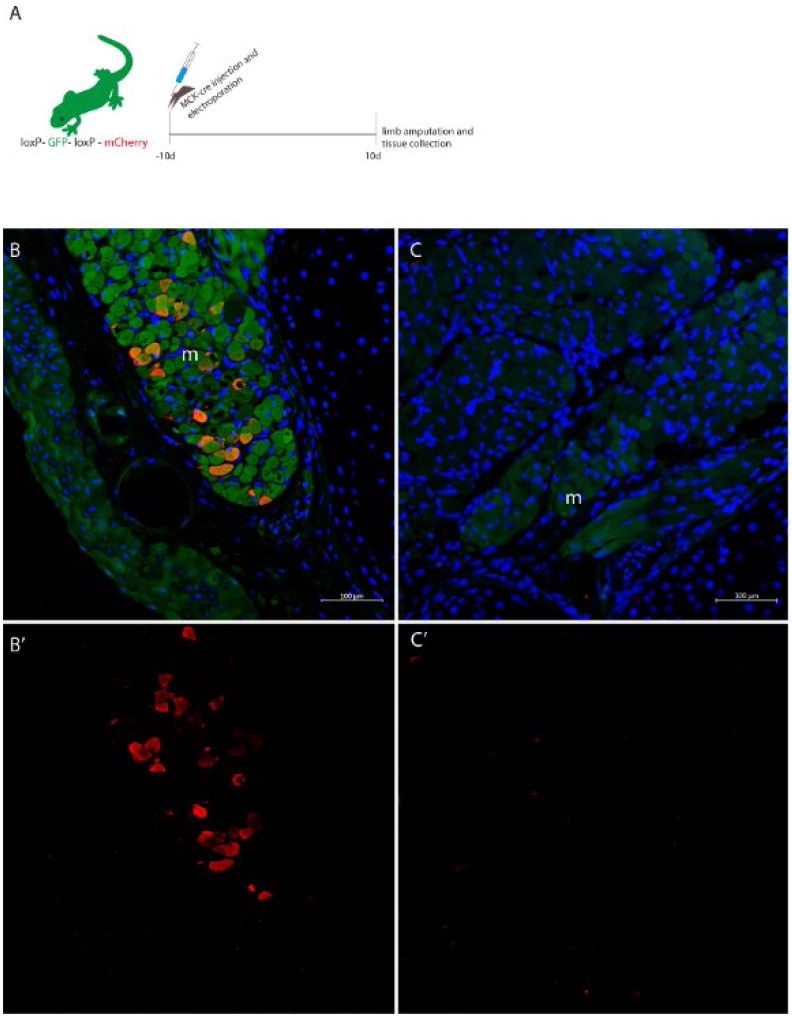
(**A**) Induction of MCK-Cre by electroporation in loxP-GFP-loxP-mCherry newt limbs. Focal electroporation leads to excision of GFP and expression of mCherry in muscle fibers. (**B**) Cross section of the newt limb 10d post-electroporation. (**B’**) The panel shows mCherry expression in muscle bundles. (**C**) Cross section of contralateral non-electroporated control limb. (**C’**) The panel shows the absence of muscle-specific mCherry expression. M-muscle bundles.

**Supplement Figure 3.**
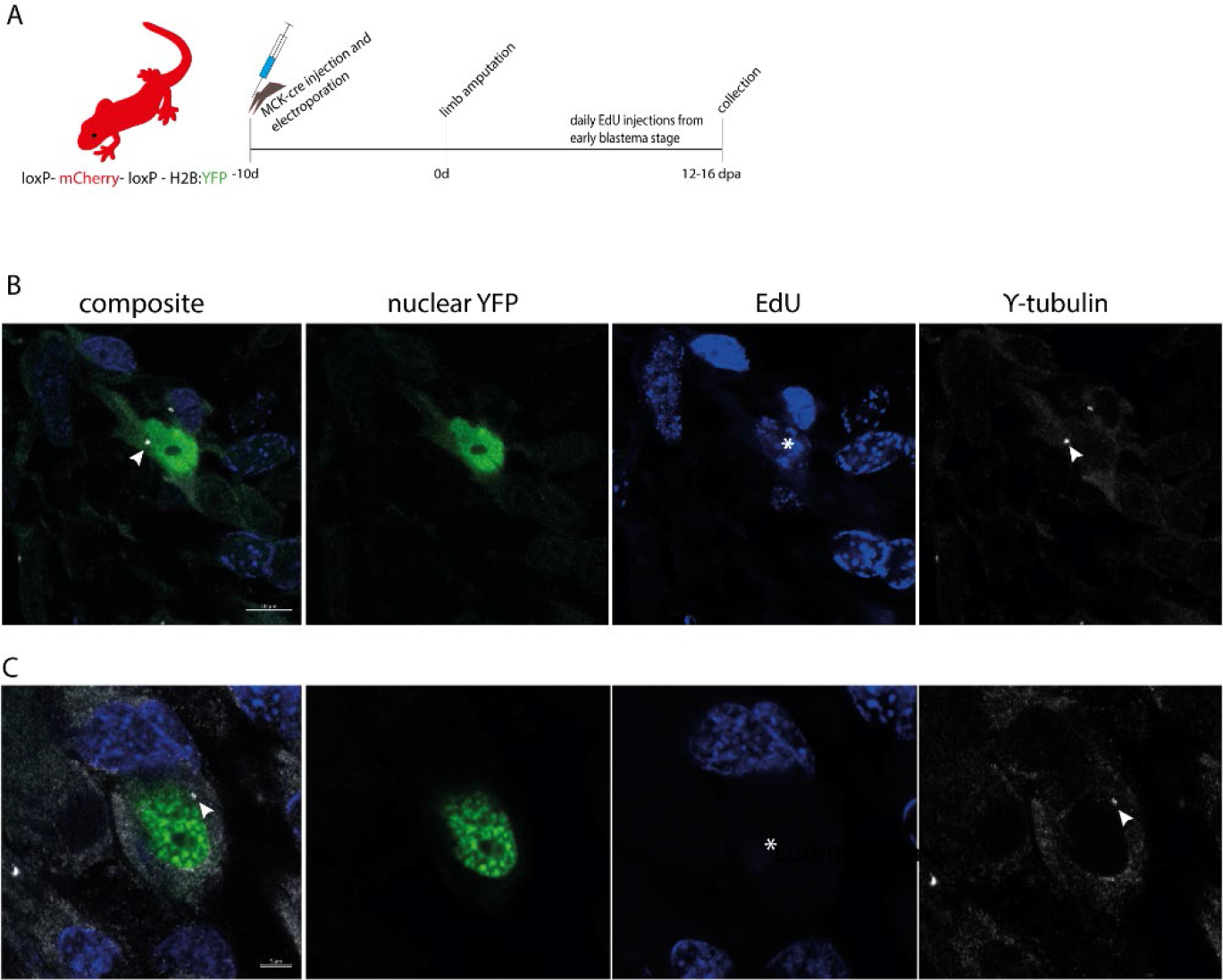
(**A**) Schematic illustration of MCK-Cre induction to induce nYFP expression in muscle fibers. (**B**) A representative image showing dedifferentiated mononucleate cells in limb sections. The cell has entered S phase as indexed by EdU (white asterisk). The centrosome within the cell is arrowed white. (**C**) A dedifferentiated mononucleate cell without S phase entry (asterisk) showing presence of centrosome (arrowed white). Scale bars, 5 µm.

**Supplementary figure 4.**
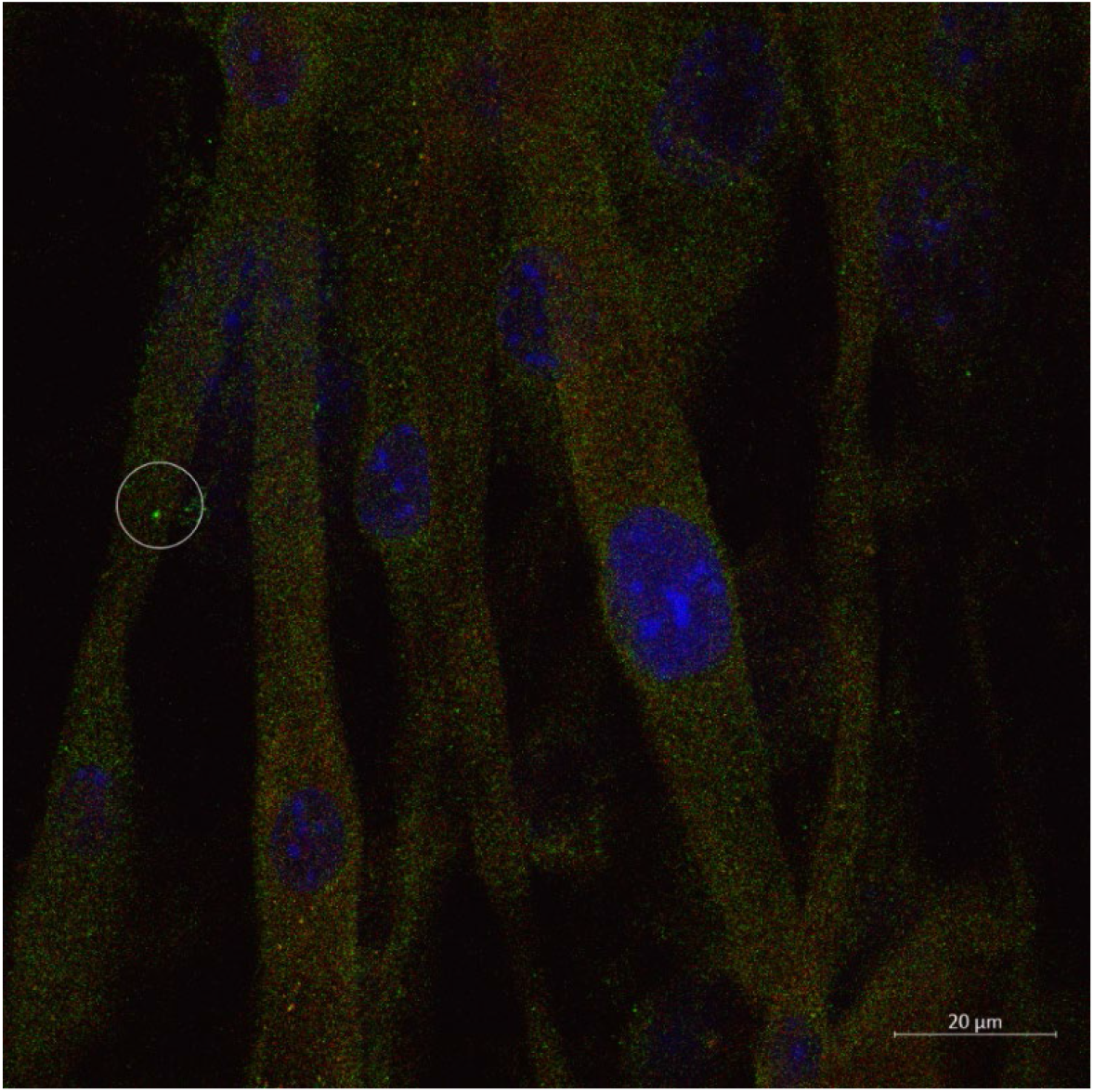
Representative image showing depletion of residual centrosomes from myotubes after centrinone treatment. The myoblasts were treated with 5 µM centrinone for 4 days during myoblast culture and 2d during differentiation. The cultures were fixed and stained with antibody to mCherry and Ƴ-tubulin. Ƴ-tubulin staining shows lack of centrosomes in 99.4% myotubes. Residual centrosomes (circled white) remaining among myotubes is estimated at 0.6% (N = 181 myofiber nuclei from 15 image fields).

**Supplementary figure 5.**
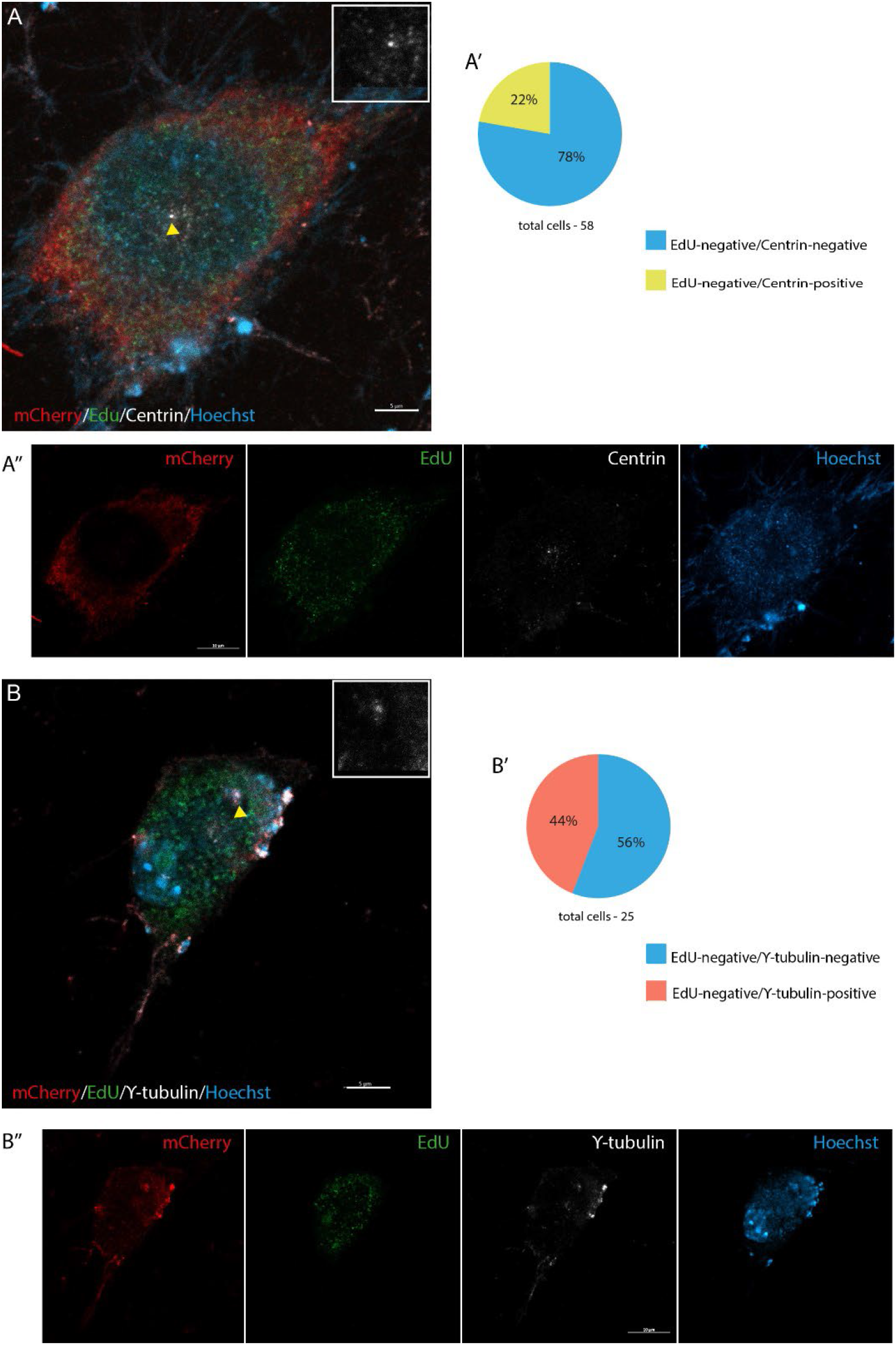
Relationship to cell cycle reentry and regain of centrosomes in myotube-derived mononucleate cells. (**A**) Representative image showing mCherry+/EdU-cell with immunoreactivity for centrin (arrowed yellow). (**B**) Representative image showing mCherry+/EdU-cell with reactivity for Ƴ-tubulin (arrowed yellow). Inset, higher magnification of a centrosome. (**B’**) Distribution of centrosome markers among myotube-derived mononucleate cells. Scale bars, 5 µm.

**Supplementary figure 6.**
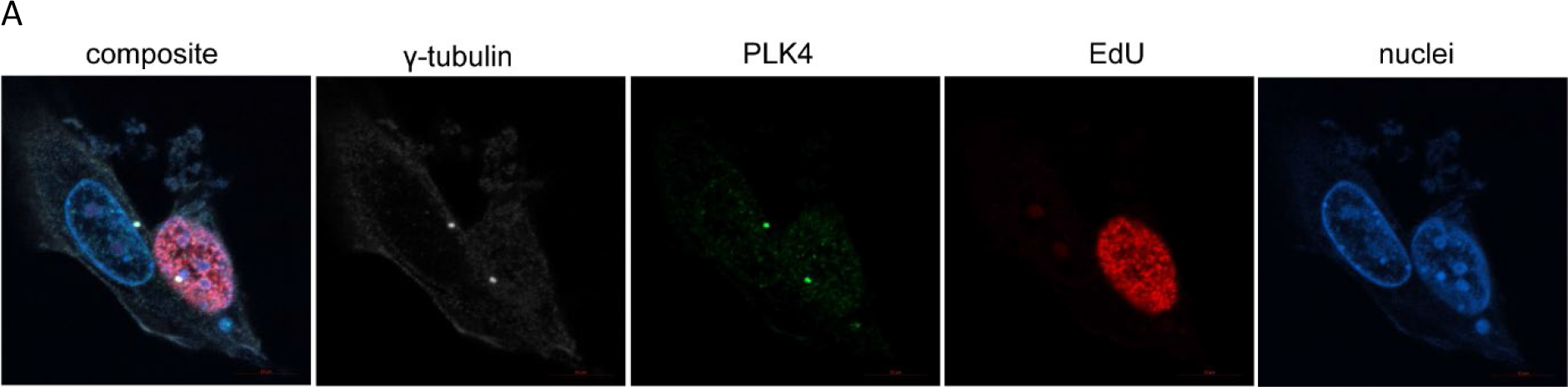
(A) PLK4 localization is not cell cycle dependent. Cultured pmi28 myoblasts showing expression of PLK4 protein and its colocalization with Ƴ-tubulin in both an S-phase and nonproliferating cell.

**Supplementary figure 7.**
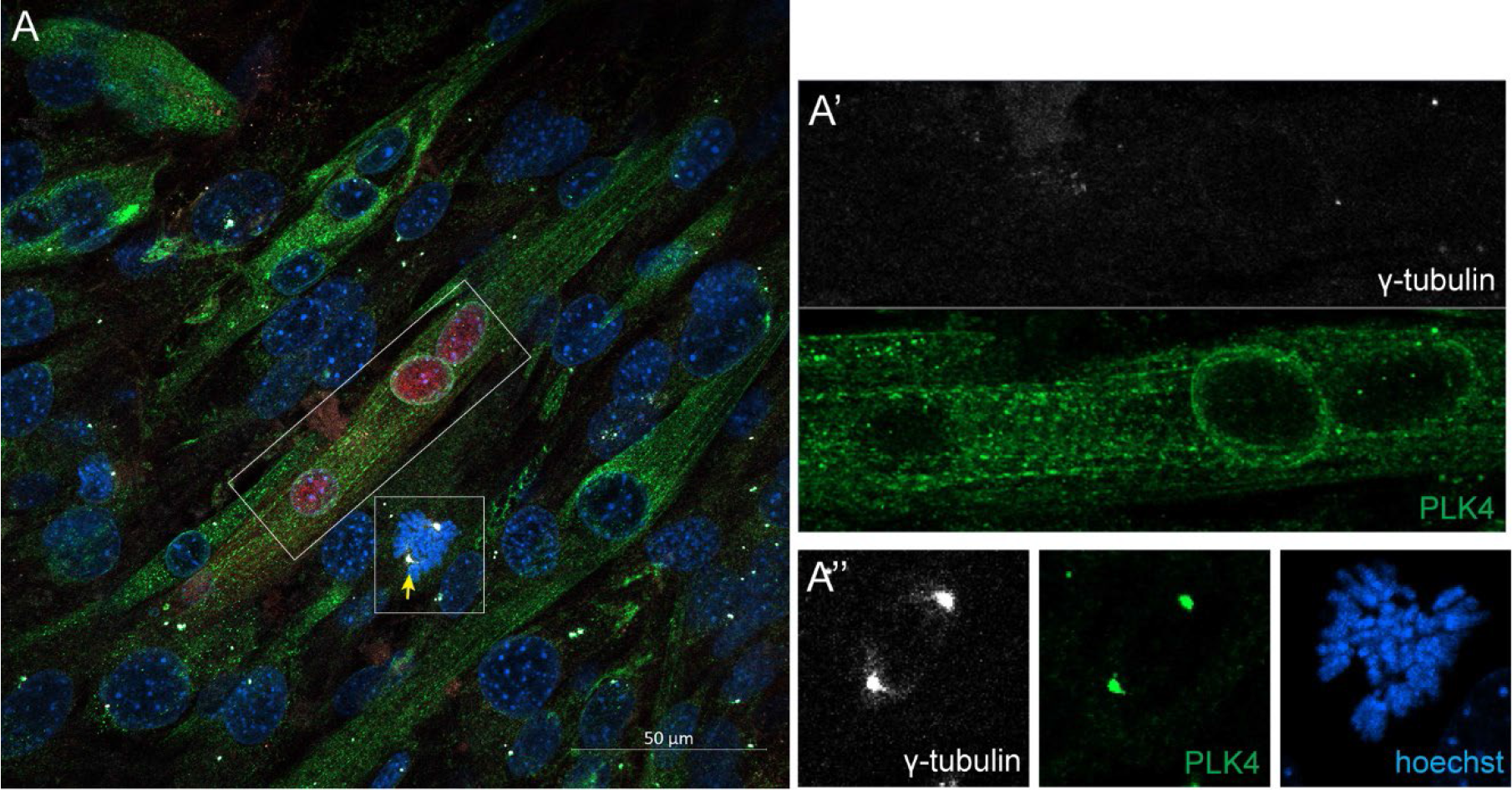
**(A)** Differentiated cultures of myotubes showing PLK4 distribution. A Cre-recombinase induced tdTomato+ myotube (rectangle in (A) showing cytoplasmic redistribution and perinuclear localization of PLK4 (magnified area in **(A’)**. Note the centrosomal localization of PLK4 in residual mononucleate cells including a mitotic cell (arrowed yellow in the box; magnified in **A”**).

**Supplementary figure 8.**
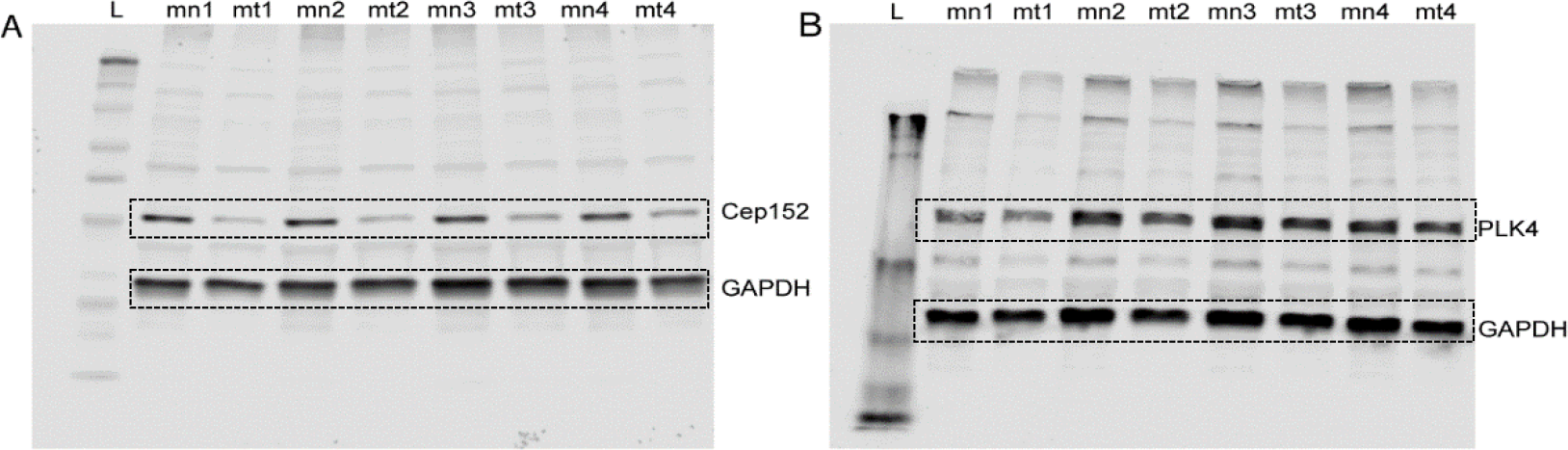
CEP 152 and PLK4 expression in cultured mononucleate and differentiated muscle cells. Western blot showing the protein levels of Cep152 (A) and PLK4 (B) in purified myotubes (mt) and mononucleate (mn) cells. The assays were represented in quadruplicate samples with GAPDH as a control. (Related to Figure 5).

**Supplementary figure 9.**
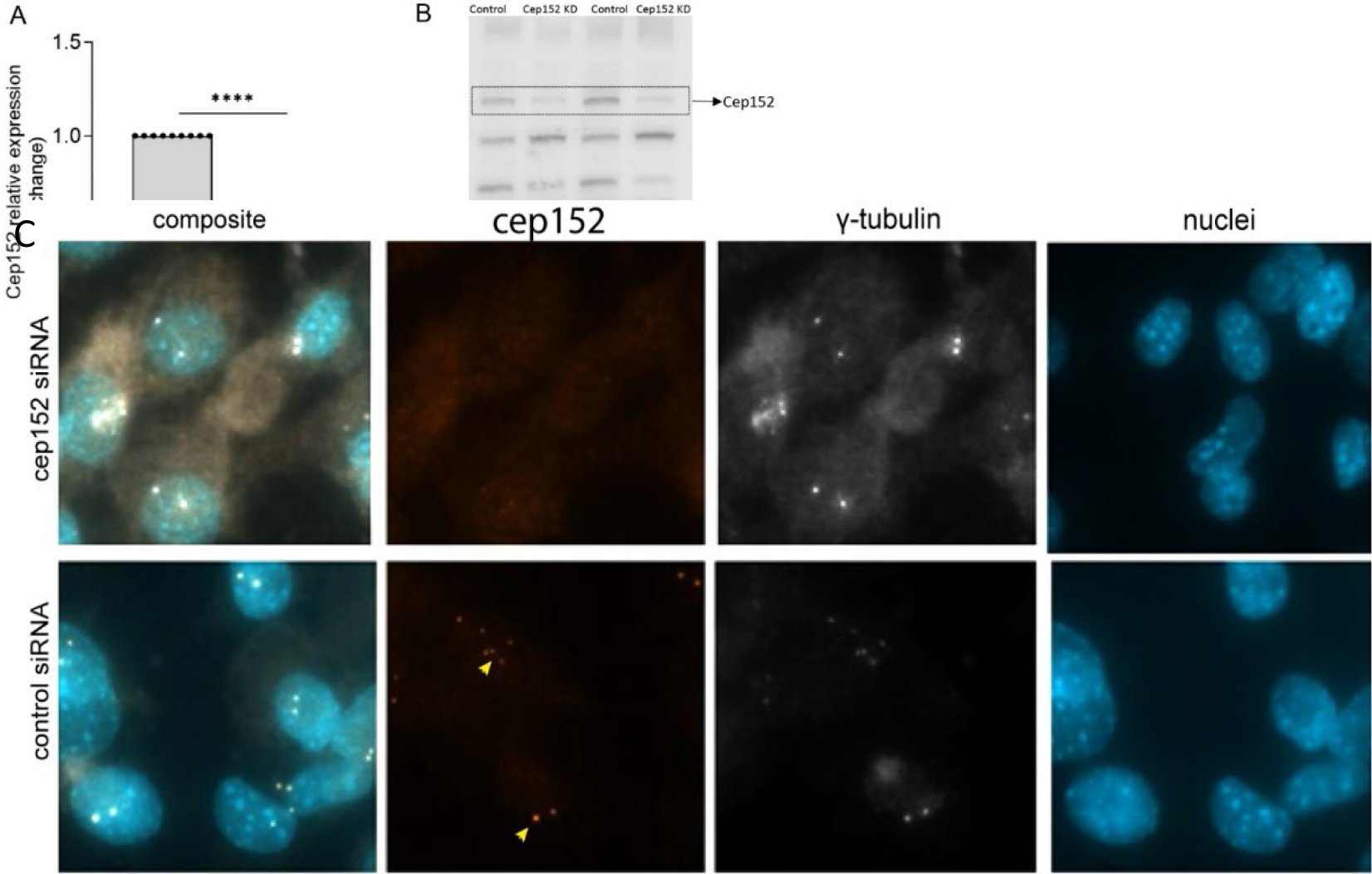
Cep152 RNAi treatment in C2C12 myoblast cells. (A) Relative expression of cep152 mRNA level in cultured myoblast cells after cep152 RNAi. P< 0.0001. (B) Western blot showing down regulation of cep152 protein expression in cultured myoblasts after Cep152 RNAi. Panels denote two independent experiments. (C) DsiRNA-mediated knockdown of Cep152 mRNA results in depletion of cep152 protein in centrosomes (upper panel). Control DsiRNA treatment (lower panel) had no effect on cep152 protein associated with centrosomes. Centrosomal ϒ-tubulin levels remain unchanged in both cases.

**Supplementary figure 10.**
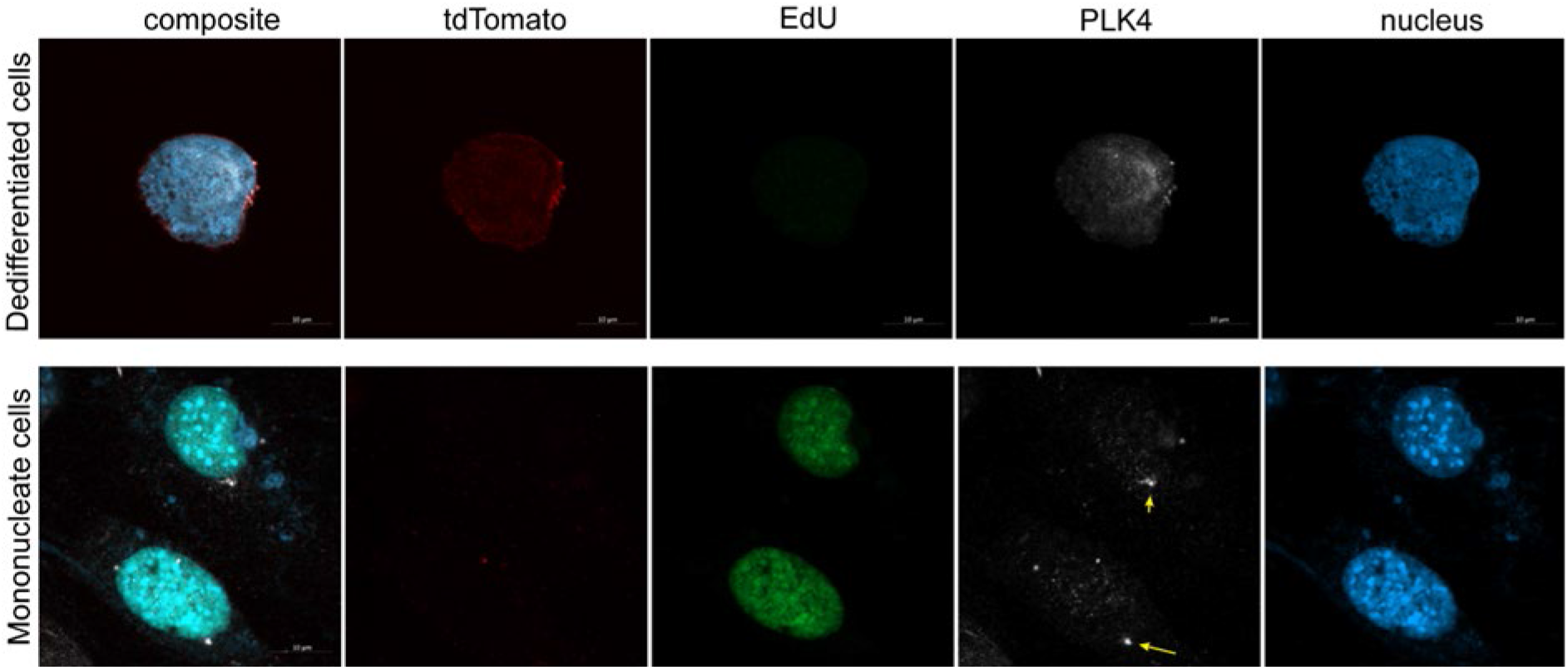
Cellularization in the absence of p53 inhibition. A representative tdTomato+ mononucleate cell showing lack of S phase entry and centrosomal PLK4 (upper panel). Residual mononucleate cells from the culture show cell cycle reentry and PLK4 expression in centrosome (lower panel).

## Notes

### Competing Interest Statement

The authors have declared no competing interest.

